# memo-eQTL: DNA methylation modulated genetic variant effect on gene transcriptional regulation

**DOI:** 10.1101/2023.05.02.539122

**Authors:** Yong Zeng, Rahi Jain, Musaddeque Ahmed, Haiyang Guo, Yuan Zhong, Wei Xu, Housheng Hansen He

**Affiliations:** Princess Margaret Cancer Centre, University Health Network, Toronto, Canada; Department of Medical Biophysics, University of Toronto, Toronto, Canada; Dalla Lana School of Public Health, University of Toronto, Toronto, ON, Canada; Department of Clinical Laboratory, the Second Hospital, Cheeloo College of Medicine, Shandong University, Jinan 250033, Shandong, China

**Keywords:** SNP, CTCF, meCpG, eQTL, memo-eQTL, Chromatin 3D structure

## Abstract

Expression quantitative trait locus (eQTL) analysis has become an important tool in understanding the link between genetic variants and gene expression, ultimately helping to bridge the gap between risk SNPs and associated diseases. Recently, we identified and validated a specific case where the methylation of a CpG site can affect the relationship between the genetic variant and gene expression. To systematically evaluate this regulatory mechanism, we developed an extended eQTL mapping method named DNA methylation modulated eQTL (memo-eQTL). We performed memo-eQTL mapping in 128 normal prostate samples and discovered 1,731 memo-eQTLs, a vast majority of which have not been reported as eQTLs. We found that the methylation of the memo-eQTL CpG sites can either enhance or insulate the interaction between SNP and Gene expression by altering CTCF-based chromatin 3D structure. This study demonstrated the prevalence of memo-eQTLs, which can enable the identification of novel causal genes for traits or diseases associated with genetic variations.

## Background

Genome-wide association studies (GWAS) have identified over 300,000 SNP-trait associations [1], but the vast majority (> 90%) of these disease-associated risk single-nucleotide polymorphisms (rSNPs) are located in non-coding regions, complicating their functional evaluation [2]. Expression quantitative trait locus (eQTL) mapping is a valuable tool to elucidate the relationship between genetic variants and transcriptional gene expression, helping to bridge the gap between rSNPs and associated diseases by identifying potential target genes [3,4]. There is significant overlap between rSNPs and eQTLs than would be expected by chance (reviewed in [5]), and these overlapped SNPs are often enriched in cis-regulatory elements (CREs) that affect transcriptional regulation [6]. Despite this, a large number of rSNPs remain untagged by any target genes through eQTL analysis.

Recently, we identified prostate cancer rSNP, rs11986220, as a novel eQTL for the oncogenic gene MYC in a subset of samples with a higher level of methylation at a CpG site located approximately 10 kilobase pairs (kbp) upstream of MYC promoter. We demonstrated that higher DNA methylation at this site prevents CTCF binding and the formation of a chromatin loop, which would allow for long-range interaction between this rSNP and MYC [7]. Unlike SNPs that are located in CTCF-binding sites and directly affect high-order chromatin architecture [8,9], this rSNP is located in an enhancer region ∼210kbp away from the CTCF binding site [7]. This study highlights a functional mechanism in which DNA methylation acts as a moderator to regulate the relationship between genetic variant and gene expression by affecting CTCF-based 3D chromatin architecture.

To systematically evaluate the DNA methylation modulated relationship between genetic variant and gene expression, we proposed a new eQTL analysis method called DNA methylation modulated eQTL (memo-eQTL). The modulation effect was statistically characterized as the interaction between the SNP and methylated CpG site (SNP × meCpG), and its significance was determined by comparing the multiple regression models with and without the interaction [10,11]. Analysis in 128 normal prostate tissue samples identified 1,732 memo-eQTLs, of which 1,720 were novel eQTLs. This method has the potential to identify a new category of eQTLs that undergo DNA methylation modulation.

## Results

### Identification of meCpG sites associated with CTCF occupancy

Our previous study had shown a meCpG can modulate the interplay between genetic variant and gene expression by altering CTCF binding [7]. Based on this, we hypothesized that other meCpGs sites, whose methylation levels are associated with CTCF occupancy, could have similar effects. To investigate this, we analyzed the correlation of 741,933 meCpG-CTCF pairs based on 26 human cell lines or tissues, including the prostate gland, using matched whole-genome bisulfite sequencing (WGBS) and CTCF chromatin immunoprecipitation sequencing (ChIP-seq) data from ENCODE (**Fig. S1A, B**) [12]. We found that 45,381 meCpG-CTCF pairs showed significantly negative correlation, while only 1,307 pairs displayed a positive correlation (**Fig. S1C** and **Supplementary Table 1**).

Although a single CTCF-binding site might contain multiple meCpG sites (**Fig. S1D**), we only identified meCpG-CTCF pairs showing opposite correlations in 23 CTCF-binding sites, suggesting that meCpG sites located in the same CTCF-binding site would have a similar relationship with CTCF occupancy. Thus, after removing those 23 CTCF-binding sites, we selected the most significantly correlated meCpG-CTCF pair for each CTCF binding site, resulting in 16,124 negatively correlated and 763 positively correlated meCpG-CTCF pairs, respectively (**Fig. 1A, B**). This is consistent with previous findings that methylation levels of meCpGs are primarily negatively associated with CTCF occupancy [13,14].

**Figure 1:**
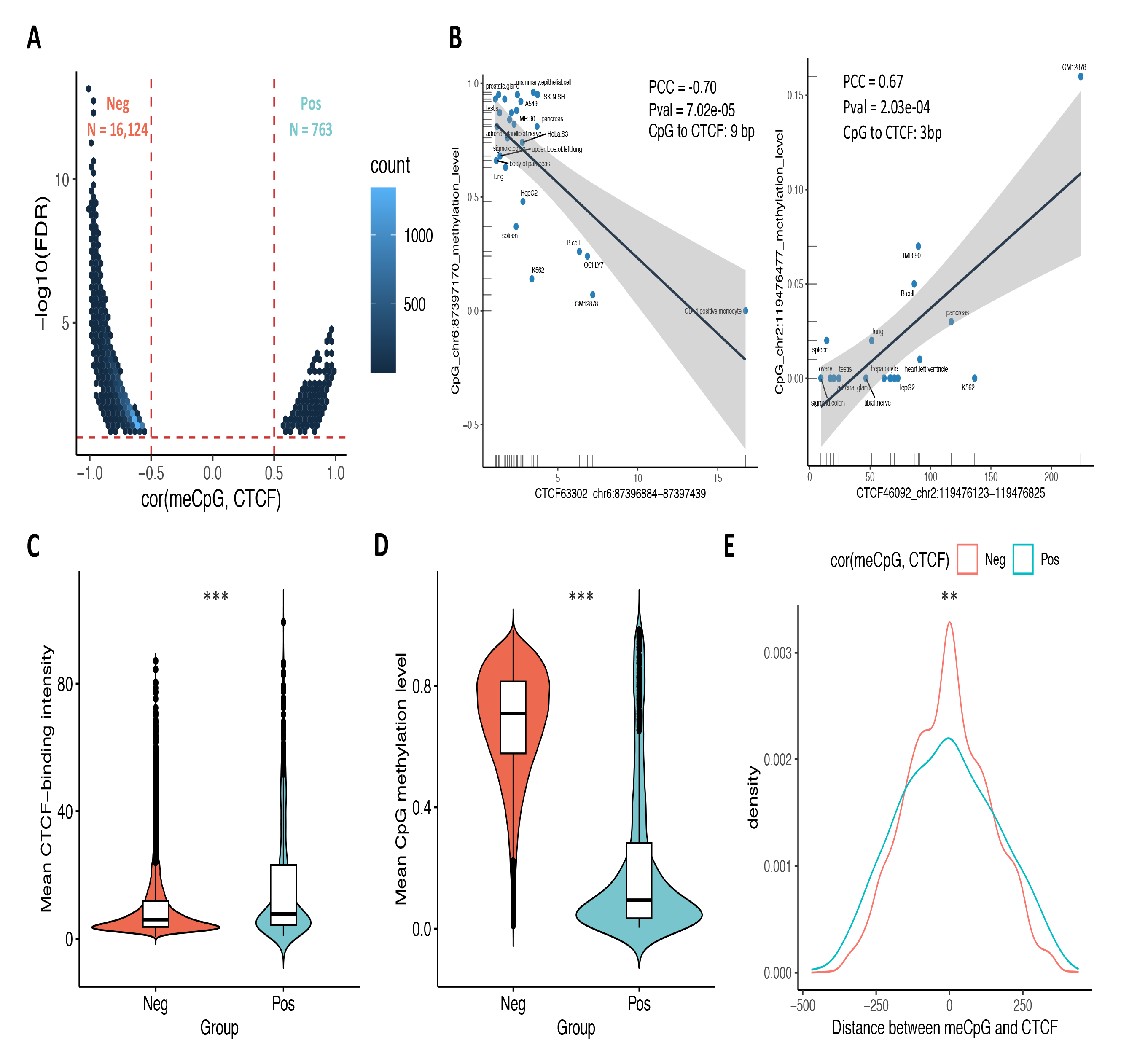
Correlation between meCpG and CTCF binding. **A)** The correlation coefficients and statistical significance for the most significantly correlated meCpG and CTCF per each CTCF binding site. Neg and Pos refer to negatively correlated and positively correlated meCpG-CTCF pairs, respectively. **B)** Examples of negatively and positively correlated meCpG-CTCF pair across 26 ENCODE samples (PCC: pearson correlation coefficient; Pval: p value; CpG to CTCF: the distance from the meCpG site to the center of the CTCF-binding site). Comparisons of the average CTCF binding intensity (**C**) and average CpG methylation levels (**D**) between Neg and Pos groups (Wilcoxon rank-sum two-sided test: mean CTCF intensity: p < 2.20 × 10^-16^; mean CpG methylation level: p < 2.20 × 10^-16^). **E)** Comparison of the distances between the meCpG site and the center of CTCF-binding site for Neg and Pos groups (Kolmogorov-Smirnov test: p = 2.08 × 10^-3^). **p < 0.01; ***p < 0.001.

Our analysis revealed that meCpG-CTCF pairs showing negative correlation tend to have higher CpG methylation levels (**Fig. 1C**) and lower CTCF occupancy compared to those showing positive correlation (**Fig. 1D**). Additionally, the meCpG sites negatively associated with CTCF binding are more likely to be located closer to the center of the corresponding CTCF-binding sites (**Fig. 1E** and **Fig. S1E**). In total, we obtained 16,887 meCpG-CTCF pairs that were significantly associated for further examination of the modulation effect on eQTL (**Supplementary Table 1**).

### memo-eQTL mapping reveals hidden relationship between SNP and gene

We proposed an extended eQTL method called memo-eQTL mapping to systematically assess the modulation effects of these meCpGs, This method characterizes the modulation effect as the interaction between SNP and meCpG (SNP × meCpG) via a moderate model (**M3**). Subsequently, it requires comparisons against the covariate model (**M2**) and the standard eQTL model (**M1**) to determine the statistical significance of the modulation effect (**Fig. 2A** and **Methods**).

**Figure 2:**
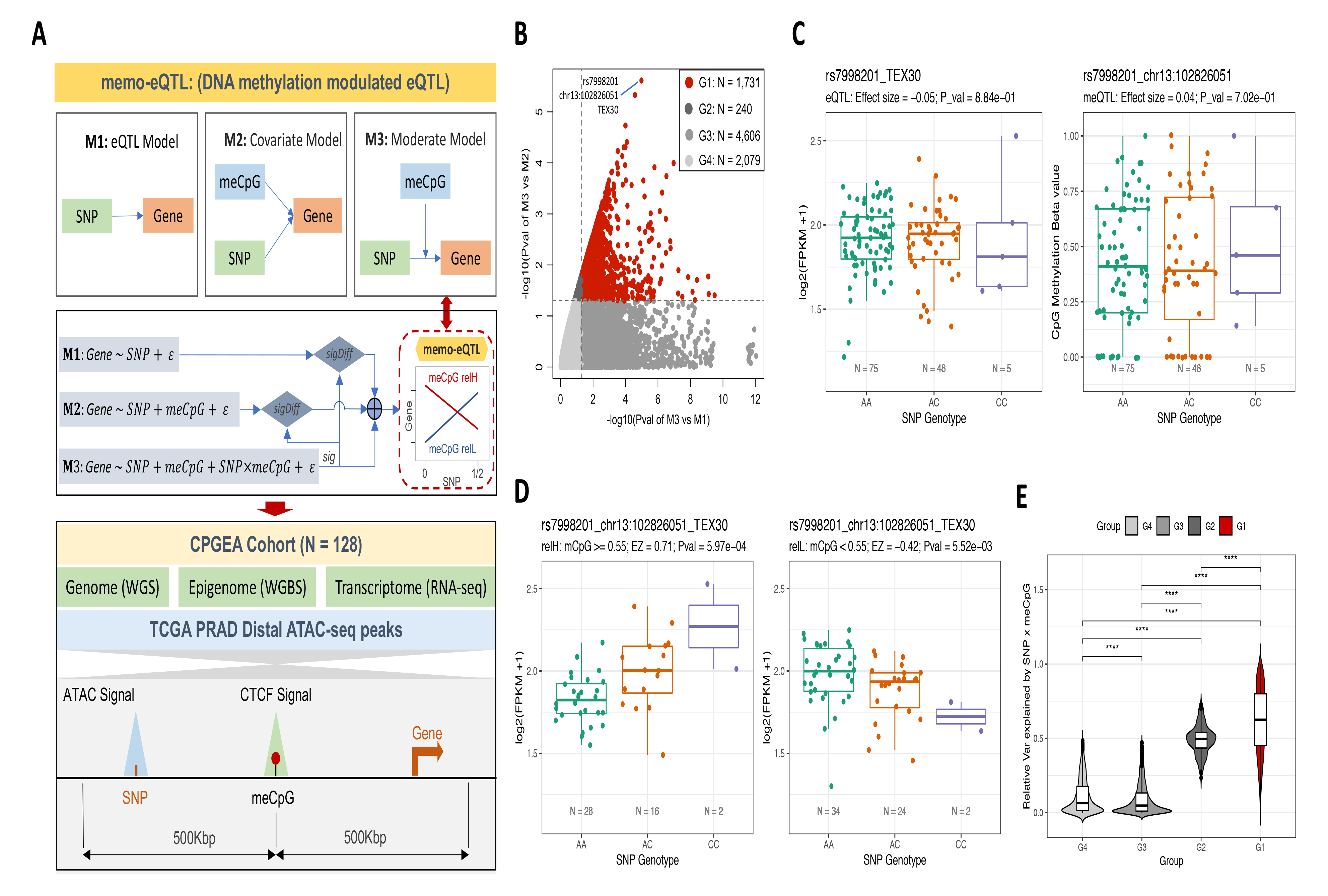
Mapping and characteristics of memo-eQTLs. **A)** The framework of memo-eQTL mapping method and its implementation in the CPGEA cohort (sig: significant; sigDiff: significantly different; relH and relL refer to the subsamples with relatively high and low methylation levels at corresponding meCpG site, respectively). **B**) Four different groups of SNP-meCpG-Gene combinations based on comparisons of M3 versus M1 and M3 versus M2 after requiring that M3 be significant. Note that combinations belonging to the group 1 (G1) are considered as memo-eQTLs. **C)** Canonical eQTL (left) and meQTL (right) analysis for SNP rs7998201 with gene TEX30 and CpG site at chr13:102826051, respectively. **D)** The visualization of selected memo-eQTL, depicting the relationship between rs7998201 and TEX30 in subsamples with relatively high (relH: Beta >= 0.55) and low (relL: Beta < 0.55) methylation levels at chr13:102826051. **E)** The comparisons of the relative variance of gene expression can be explained by SNP × meCpG across groups G1-4 (Wilcoxon rank-sum two-sided test: ****p < 0.0001).

We conducted the memo-eQTL mapping in the Chinese Prostate Cancer Genome and Epigenome Atlas (CPGEA) cohort, which included matched whole-genome sequencing (WGS), RNA sequencing (RNA-seq) and WGBS data for 128 benign prostate samples [15] (**Fig. 2A**). Specifically, we pruned SNPs in high linkage disequilibrium (LD) and focused on 19,895 SNPs located in 14,374 ATAC-seq distal peak regions that were identified in prostate cancer [16] (**Fig. S2A** and **Methods**). For meCpG sites, we pinpointed 5,525 sites with variable methylation levels (**Fig. S2B** and **Methods**), which were significantly correlated with CTCF-binding as reported in **Fig.1A**. Among the potential target genes, we preserved 14,520 protein-coding and lincRNA genes after filtering out lowly expressed ones (Median FPKM < 1, **Fig. S1C**). Lastly, we conducted memo-eQTL analysis for 90,959 SNP-meCpG-Gene combinations, where the linear distance between the SNP and the Gene was up to 1 million base pairs. Importantly, we required the meCpG site to be located in between the paired SNP and Gene to simplify the possible modulating mechanisms (**Fig. 2A** and **Methods**).

In total, we identified 1,731 memo-eQTLs, which not only displayed a statistically significant SNP × meCpG interaction (**M3** versus **M2**) but also showed significant improvement over the canonical eQTL models (**M3** versus **M1**) (**Fig. 2B and Supplementary Table 2**). Most of SNP and gene pairs (1,720/1,731) had not been reported by GTEx eQTL analysis in the prostate (dbGaP Accession phs000424.v8.p2), and only 160 and 134 of these memo-eQTLs were also detected as eQTLs and methylation quantitative trait loci (meQTLs) in the CPGEA cohort, respectively. For instance, the SNP rs7998201 is neither an eQTL for TEX30 nor a meQTL for the CpG site at chr13:102826051. However, it is significantly associated with TEX30 expression levels in the subsamples with either high or low methylation levels at chr13:102826051 (**Fig. 2B-D** and **Methods**). It suggests that this meCpG site can modulate the relationship between the rs7998201 and TEX30 expression. Additionally, we found the SNP × meCpG can explain the highest relative percentage of gene expression variance for memo-eQTLs compared to other non-significant groups (**Fig. 2B, E**). In contrast, the SNP or meCpG alone tends to explain less gene expression variance for memo-eQTLs compared to other groups (**Fig. 2B** and **Fig. S2D**).

In conclusion, these results suggest that memo-eQTL mapping can complement canonical eQTL and meQTL analyses and specifically uncover the hidden relationship between genetic variant and gene expression modulated by DNA methylation.

### The characteristics of meCpGs, genes and SNPs involved for memo-eQTLs

There are 808, 1,197 and 1,344 unique meCpGs, genes and SNPs involved for the 1,731 memo-eQTLs, which are termed as eCpGs, eGenes and eSNPs hereinafter, respectively (**Fig. 3A**). Notably, an eCpG can modulate the relationships of up to 42 pairs of eSNP and eGene, and an eGene can also be associated with as many as 11 combinations of eCpG and eSNP, indicating the possible dominance or additive modulation effects of eCpGs (**Fig. 3A**). To explore the biological processes and functions regulated by memo-QTLs, we performed Gene Ontology (GO) enrichment analysis for all eGenes and found that they are enriched on chr6p21 and over-represented in several immune related Kyoto Encyclopedia of Genes and Genomes (KEGG) pathways (**Fig. 3B, C**), especially in the antigen processing and presentation pathway, which plays a key role in adaptive immunity [17]. However, due to the high linkage disequilibrium in the MHC region on chr6p21 [18], further investigations are needed to determine which eGene is responsible for the associated eSNPs.

**Figure 3:**
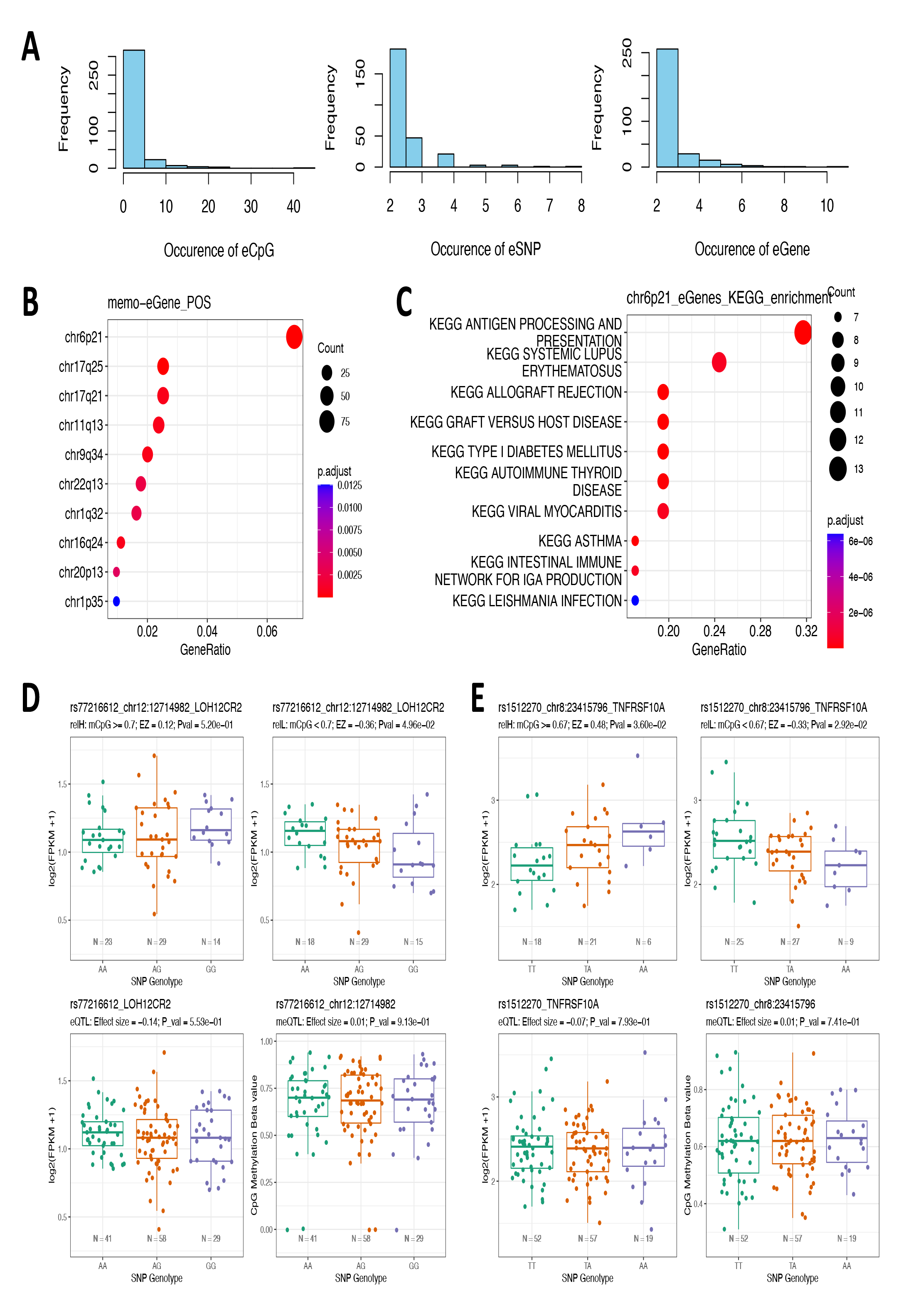
Characteristics eCpG, eGene and eSNP for memo-eQTLs. **A)** The occurrence of eCpG (left), eSNP (middle) and eGene (right) in 1,731 memo-eQTLs. **B)** Enrichment of eGenes in various chromosome regions. **C)** Enriched KEGG pathways for eGenes located in chr6p21. **D, E)** The visualization of memo-eQTLs, eQTLs and meQTLs for prostate cancer risk SNP rs77216612 and its high LD eSNP rs1512270.

Of the 1,344 unique eSNPs, 65 have been previously reported to be associated with 41 types of traits or diseases, such as hypertension, diabetes, basal cell carcinoma, and prostate cancer, in GWAS catalog studies (**Supplementary Table 3**) [1]. An additional 265 eSNPs are in high LD with significant risk SNPs in GWAS catalog (**Methods**). Notably, we identified the prostate cancer risk SNP rs77216612 as an eSNP, which is associated with the expression of two long noncoding RNAs (lncRNAs), LOH12CR2 and RP11-253I19.3, located in a tumor suppressor locus 12p12-13 [19] and modulated by an eCpG site at chr12:12714982 (**Fig. 3D** and **Fig. S3A**). In addition, two other eSNPs in high LD with this rSNP, rs17194607 and rs1512270, are associated with the expression of C6orf136 and TNFRSF10A, which are modulated by eCpGs at chr6:30238902 and chr8:23415796, respectively (**Fig. 3E** and **Fig. S3B**). It is worth noting that none of these 3 eSNPs were detected as canonical eQTLs or meQTLs (**Fig. 3E** and **Fig. S3B**), and the transcriptional regulation mechanisms of TNFRSF10A, a critical cell surface receptor that binds to tumor necrosis factor-related apoptosis-inducing ligand and mediates the extrinsic apoptosis pathway, are largely unknown [20]. These results suggest that memo-eQTLs can complement eQTLs and help interpret the associations between SNPs and traits or diseases detected by GWAS.

### eCpG-CTCF based chromatin loop can either insulate or enhance the regulatory interaction between eSNP and eGene

In our previous study, we demonstrated that a high CpG methylation level can prevent CTCF binding and formation of a 3D chromatin loop, enabling cross-talk between a SNP and its target genes, while a low CpG methylation level is associated with more CTCF binding and formation of the 3D loop, which acts as an insulator blocking the interplay between the SNP and gene [7]. To systematically assess the underlying regulatory mechanism for memo-eQTLs, we categorized them into four groups based on whether significant associations between eSNPs and eGenes were observed in subsamples with high and low eCpG methylation levels: sigHigh, sigLow, sigBoth, and sigNone memo-eQTLs (**Methods**). To simplify the categorization, we excluded the 20 memo-eQTLs with meCpG sites positively correlated with CTCF binding, resulting in 627 sigHigh, 621 sigLow, 203 sigBoth and 260 sigNone memo-eQTLs (**Fig. 4A**).

**Figure 4:**
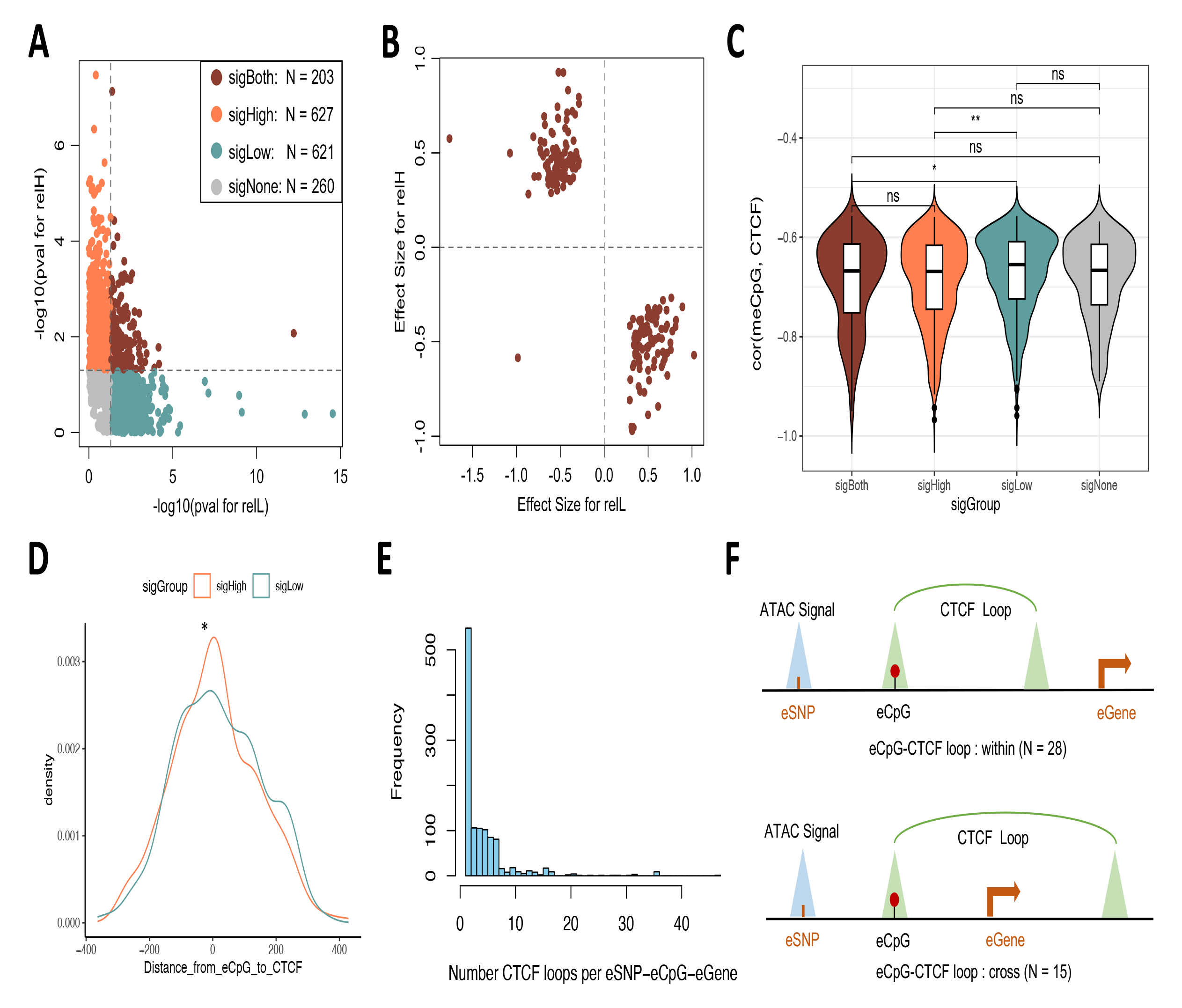
Mechanisms investigation for memo-eQTLs. **A)** Stratified four subgroups of memo-eQTLs based on the p values of **M1** models for relatively high (relH) and low (relL) subsamples of the eCpG site, 0.05 was used as the significance cutoff. **B)** The effect size and direction of eSNPs on eGenes in eCpG relH and relL subsamples for the sigBoth memo-eQTLs. **C)** Comparisons of the correlation coefficients for meCpG-CTCF pairs across four memo-eQTL groups (Wilcoxon rank-sum two-sided test: ns: not significant; *p < 0.05; **p < 0.01). **D)** Distribution and comparison of distance between eCpG and the center of the associated CTCF-binding site for sigHigh and sigLow memo-eQTLs (Kolmogorov-Smirnov test: p = 4.79 × 10^-2^). **E)** The number of RWPE1 ChIA-PET data derived CTCF loops that overlapped with the eSNP-eCpG-eGene loci. **F)** The illustration plot of the overlapping patterns between eSNP-eCpG-eGene loci and eCpG-CTCF loops. The numbers represent results using RWPE1 ChIA-PET data.

Interestingly, we observed the eSNPs tend to have opposite relationships with eGenes in subsamples with relatively high and low eCpG methylation levels, particularly for the sigBoth memo-eQTLs (**Fig. 4B** and **Fig. S4A**). This finding suggests that the meCpG modulation effect not only determines the presence of the interaction between genetic variant and gene expression but also changes the orientation of their cross-talk. When delving into the distances among eSNP, eCpG and eGene across four types of memo-eQTLs, we found no significant differences in the pairwise linear distances (**Fig. S4B** and **Supplementary Table 4**). Moreover, the correlations between meCpG and CTCF binding were similar overall, except for the sigHigh eCpGs, which showed a slightly stronger negative correlation with CTCF occupancy (**Fig. 4C**), as well as being located closer to the center of the CTCF-binding site (**Fig. 4D**). Together, these results suggest that eCpG mainly modulates the interplay between eSNP and eGene by affecting CTCF-based chromatin organization to alter their spatial distance.

To validate this, we employed the CTCF chromatin interaction analysis with paired-end-tag sequencing (ChIA-PET) data from the prostate epithelial cell line RWPE-1 [21], and focused on the 5,068 strongest CTCF-based chromatin loops (#PET reads >= 15). We found that the eSNP-eCpG-eGene loci of 1,155 memo-eQTLs overlapped with CTCF loops, with 80% of these regions intersecting with more than one loop (**Fig. 4E**). We also identified 111 eCpGs sites located in the anchor sites of 134 CTCF loops, which allowed us to directly assess their modulation effects through 3D structure alteration. To simplify the analysis, we focused on 97 eCpGs that overlapped with a single CTCF loop anchor site, which we termed as eCpG-CTCF loop. There were 61 eCpG-CTCF loops fully embedded in corresponding eSNP-eCpG-eGene loci, while the remaining 36 eCpG-CTCF loops partially intersected with the eSNP-eCpG-eGene loci. These two groups were termed as Within and Cross eCpG-CTCF loops, respectively, as shown in **Fig. 4F**. In particular, we identified 15 sigHigh memo-eQTLs with Cross eCpG-CTCF loops (**Fig. S4C and Supplementary Table 4**), suggesting the corresponding eCpGs might block the formation of CTCF loops, which could act as insulators for the interplay between eSNP and eGene. In contrast, we also identified 24 sigLow memo-eQTLs with Within eCpG-CTCF loops (**Fig. S4C and Supplementary Table 4**), implying the corresponding eCpGs might promote the formation of CTCF loops, which could enhance the interaction between the eSNP and eGene by bringing them physically closer. Similar results were observed when we examined the HiChIP data derived chromatin 3D loops in prostate cancer cell line VCaP (**Fig. S4D, E**). In summary, we found that meCpG modulated CTCF-based chromatin 3D organization could either insulate or enhance the cross-talk between genetic variant and gene expression. However, further investigations are needed to fully uncover these complicated mechanisms.

## Discussion

In contrast to canonical eQTL or meQTL that examine the association between gene expression and genetic variant or DNA methylation alone, our previous research revealed that DNA methylation can modulate the interplay between genetic variants and gene expression by dichotomizing the population into high and low methylation groups based on a specific CpG site methylation levels [7]. To evaluate this type of modulation effect more comprehensively, we developed a more sophisticated method called memo-eQTL. This approach not only incorporates the genetic variant and DNA methylation but also their interaction into a multiple regression model. The statistical significance of the DNA methylation modulation effect is determined by comparing this model with and without the interaction [10,11]. We used the original continuous methylation levels instead of dichotomizing it as a categorical variable in our previous study, allowing us to explore the effects of genotypes at different methylation levels. To better visualize the difference in the effect of SNP on gene expression at relatively high and low CpG methylation levels, we searched for the optimal separation reflecting more striking modulation effects.

We first examined the correlation between CTCF occupancy and CpG methylation across 26 cell types and tissues. Although 97% of significant pairs are negatively correlated, a small subset of pairs exhibit positive relationships, which has also been observed in previous studies [13]. These results confirmed the overall inverse relationship between the CpG methylation and CTCF binding. However, the presence of this subset of positively correlated pairs suggests that additional factors may be involved, such as cofactors that interact with CTCF and selectively affect the binding site methylation status [22,23]. About 1% memo-eQTLs are also engaged with positively correlated meCpG-CTCF pairs, which provide an alternative meCpG modulation model for the interplay between the eSNP and eGene.

Since all the rest of the eCpGs are negatively associated with CTCF binding, we assumed that the sigHigh memo-eQTLs would preferentially partially intersect with CTCF looping, as a result, inhibiting the CTCF-based loop formation that can blockade the interplay between eSNP and eGene. Conversely, sigLow memo-eQTLs would be more likely fully located in the CTCF loop, allowing for increased eSNP and eGene interactions since the loop would bring them physically closer. However, we did not observe statistically differences between sigHigh and sigLow memo-eQTLs regarding the overlapping pattern with eCpG-CTCF loops (**Fig. S4C, E**). Moreover, there is no spatial pattern preference for sigBoth and sigNone either. One possible explanation is that the vast majority of memo-eQTL eSNP-eCpG-eGene ranges overlapped with multiple CTCF loops, making it challenging to discriminate the 3D structure differences among different groups of memo-eQTLs. Further investigations are needed to fully uncover these complicated mechanisms.

## Conclusions

The memo-eQTL method provides a valuable tool for identifying DNA methylation modulated eQTLs that are often missed by canonical eQTL analysis, thereby allowing for the discovery of novel genes that are associated with genetic variants and diseases. We found that DNA methylation modulated CTCF-based chromatin 3D organization can either insulate or enhance the cross-talk between genetic variant and gene expression. We believe that as more 3D chromatin data becomes available, our understanding of these regulatory mechanisms will continue to improve. Overall, our findings suggest that the memo-QTL method and the study of chromatin 3D organization can provide a complementary framework for identifying and understanding the complex regulatory processes that underlie genetic variation and gene expression.

## Methods

### Correlation between CpG methylation and CTCF binding

To examine the relationship between CpG methylation and CTCF-binding, we gathered a total of 95,887 CTCF-binding sites from 26 human cell lines or tissues (**Fig. S1A**), along with the methylation levels of 1,188,556 CpG dinucleotides located on these CTCF-binding sites, from the ENCODE portal [12]. For meCpG sites, we required at least 10 cell lines or tissues with more than 20 WGBS reads, as well as an Interquartile Range (IQR) of Beta values greater than 0. For the CTCF binding intensity, we required an IQR greater than 1 (**Fig. S1B**). Then, the Pearson Correlation Coefficient (PCC) and p value for each CpG site’s methylation levels and corresponding CTCF-binding intensity were calculated, and the p values were adjusted for multiplicity using the BenJamini-Hochberg Procedure. Lastly, a significant association between meCpG and CTCF binding was defined when the absolute PCC was greater than 0.5 and the Padj was smaller than 0.05.

### SNP-CpG-Gene combinations for memo-eQTL mapping

Using the processed WGS, WGBS and RNA-seq data for 128 benign prostate samples from the CPGEA cohort [15], we extracted genotype information for SNPs with rsIDs in dbSNP (build 151) and a minor allele frequency (MAF) greater than 0.05, and removed SNPs in high LD (squared correlation >= 0.8) using the PLINK pruning function (www.cog-genomics.org/plink/1.9/). Given the potentially functional capability, we focused on 19,895 pruned SNPs located in 14,374 ATAC-seq distal peak regions that were identified in prostate cancer [16] (**Fig. S1A**). For DNA methylation data, we focused on 5,525 meCpG sites with average methylation levels (Beta value) within the range of [0.25, 0.75] and IQRs greater than 0.1 (**Fig. S1B**), and that were also found significantly correlated with CTCF-binding intensity in **Fig.1A**. As for potential target genes, we retained 14,520 protein-coding and lincRNA genes after filtering out lowly expressed ones (Median FPKM < 1) and inverse normal transformed their expression levels (**Fig. S1B**). We then searched for all possible SNP-CpG-Gene combinations within a 1 million base pair window size. Importantly, we required the CpG site to be located in the middle of paired SNP and gene to ensure possible modulating effects (**Fig. 2A**).

### memo-eQTL mapping, assessment and grouping

Three models were built for memo-eQTL mapping: **M1**, the standard eQTL model, determines the effect of SNP on gene expression; **M2**, the covariate model, examines the additive marginal effects of SNP and DNA methylation on gene expression; and **M3**, the moderate model, characterizes the DNA methylation modulation effect as the interaction between the SNP and meCpG (SNP × meCpG) on top of **M2**. These models were built for all 90,959 SNP-meCpG-Gene combinations.

**M1**: Gene = α_1_ + β_1_ SNP + ε_1_
**M2**: Gene = α_2_ + β_21_ SNP + β_22_ meCpG + ε_2_
**M3**: Gene = α_3_ + β_31_ SNP + β_32_ meCpG + β_33_ SNP × meCpG + ε_3_

For memo-eQTLs, first, we required that **M3** is significant, and the contribution of the SNP × meCpG interaction is also significantly by comparing the **M3** and **M2 models** [10,11]. To ensure that the significant SNP × meCpG interaction was driving the presence or enhancement of the relationship between SNP and gene expression, we also required that the comparison between **M3** and **M1** models be significant (**Fig. A, B**). Specifically, the likelihood ratio test (LRT) [24] was employed to compare the two models, and a p-value threshold of 0.05 was used to determine statistical significance.

To visualize the effects of SNP × meCpG for memo-eQTL, we dichotomized the 128 samples based on the optimal threshold of meCpG levels that distinguished the **M1** models in meCpG relatively high (relH) and low (relL) subsamples the most. This threshold was searched within the range between the lower (Q1) and higher (Q3) quartiles of meCpG levels (**Fig. 2D**). Furthermore, we computed the relative explained variances of gene expression by SNP, meCpG and SNP × meCpG in **M3** across the G1-4 groups by normalizing the individually explained variance to their sum (**Fig. 2B, E** and **Fig. S2D**). Lastly, the memo-eQTLs were split into four groups (sigHigh, sigLow, sigBoth, and sigNone) based on the statistical significance of the **M1** models in relH and relL subsamples (**Fig. 4A**). Specifically, we examined whether these models were significant (p-value < 0.05) in either or both relH and relL subsamples.

### GWAS SNPs and their high LD SNPs

All significant SNP-trait associations were downloaded from the GWAS catalog [1], and only those SNPs with rsIDs were examined in our analysis. To identify SNPs in high LD, we used genotype data for the East Asian (EAS) population from the 1000 Genomes phase3 data [25]. To be specific, the PLINK v1.9 (www.cog-genomics.org/plink/1.9/) was employed to scan all SNP pairs within the query SNP’s centralized 1 million base-pair window, and SNPs with an R2 value greater than 0.8 were considered in high LD.

### eSNP, eGene and eCpG linear and spatial distance examination

To determine the linear distance between the eSNP, eCpG and eGene, we calculated the pairwise distances between any two of them. The distance to eGene was measured based on the transcription start site (TSS). All distances were measured strandless since meCpG is strandless, and negative distance indicates the location of the former element on the left side. To explore the chromatin 3D structures, we utilized CTCF ChIA-PET data from the normal prostate cell line (RWPE1) [21] and focused on the 5,068 strongest CTCF-based chromatin loops with at least 15 PET reads. Additionally, we called CTCF-associated chromatin loops from CTCF HiChIP data in VCaP cells (GSE172498) using HiCUP (v0.7.2) and hichipper (v0.7.7) pipelines. To simplify the investigation of mechanism, we focused on those eCpGs located in the anchor site of a single CTCF loop as derived above.

### Statistical analysis

The comparison of continuous variables between groups was conducted using the Wilcoxon rank-sum two-sided test or the Kolmogorov-Smirnov test, while the comparison of categorical variables was conducted using Chi-square test. All statistical significance were provided, and the results were considered significant if the p-value was less than 0.05.

## Availability of data and materials

The analyses were primarily performed using R 4.0.3 (http://CRAN.R-project.org, R Foundation, Vienna, Austria). All source data and code for this study will be deposited at GitHub as needed.

## Authors’ contributions

Designed studies: H.H.H, W.X and Y.Z

Data Analysis: Y.Z, R.J, M.A, H.G, Y.Z

Wrote first draft of manuscript: Y.Z

Revised & approved manuscript: all authors

## Acknowledgements

We like to thank the CPGEA team for sharing their processed WGS, RNA-seq and WGBS data.

## Supplementary Figures

**Supplementary Figure 1:**
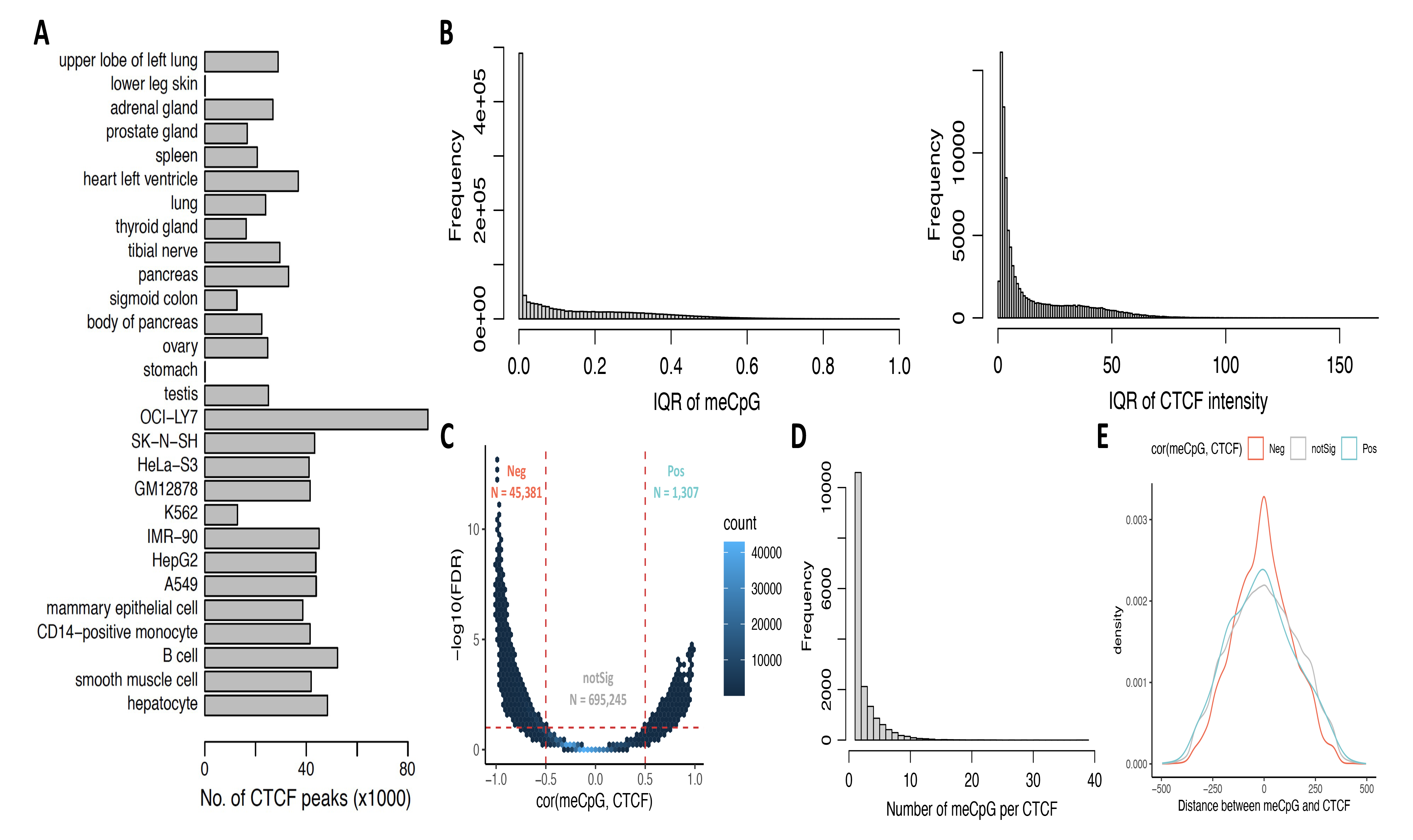
The characteristics of meCpG and CTCF binding sites. **A)** The number of CTCF peaks for 26 different tissue and cell line samples from the ENCODE. **B)** The distributions of the interquartile range (IQR) for meCpG methylation levels (left) and CTCF-binding intensity (right). **C)** Correlation coefficients and statistical significance for all meCpG-CTCF pairs. Neg, Pos and notSig refer to negatively correlated, positively correlated and not significantly correlated meCpG-CTCF pairs, respectively. **D)** Distribution of the number of meCpG sites per CTCF-binding site for significantly correlated meCpG-CTCF pairs. **E)** Comparisons of distance between meCpG site and the center of CTCF-binding site among three different groups (Kolmogorov-Smirnov test: Pos vs Neg: p = 1.69 × 10^-6^; Pos vs notSig: p = 4.58 × 10^-3^; Neg vs notSig: p < 2.20 × 10^-16^).

**Supplementary Figure 2:**
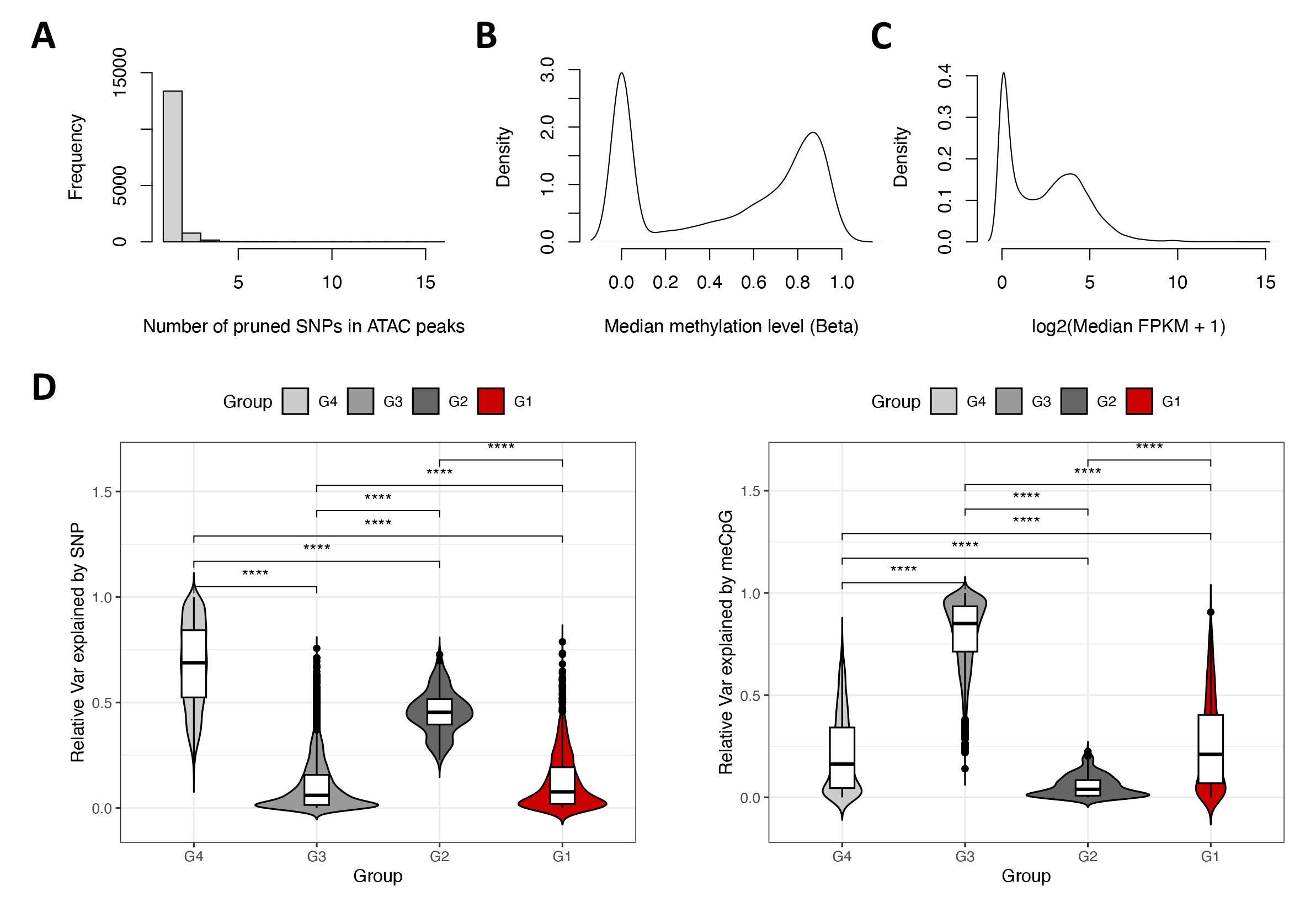
The preparation and characteristics of memo-eQTL. **A)** The distribution of the number SNPs located in ATAC peaks after filtering and pruning. **B)** The distribution of the median methylation levels for measured CpG sites across 128 samples. **C)** The distribution of the median gene expression levels for protein-coding and lincRNA genes across 128 samples. **D)** The comparisons of relative variance of gene expression can be explained by SNP (left) and meCpG (right) alone across groups G1-4 (Wilcoxon rank-sum two-sided test: ****p < 0.0001).

**Supplementary Figure 3:**
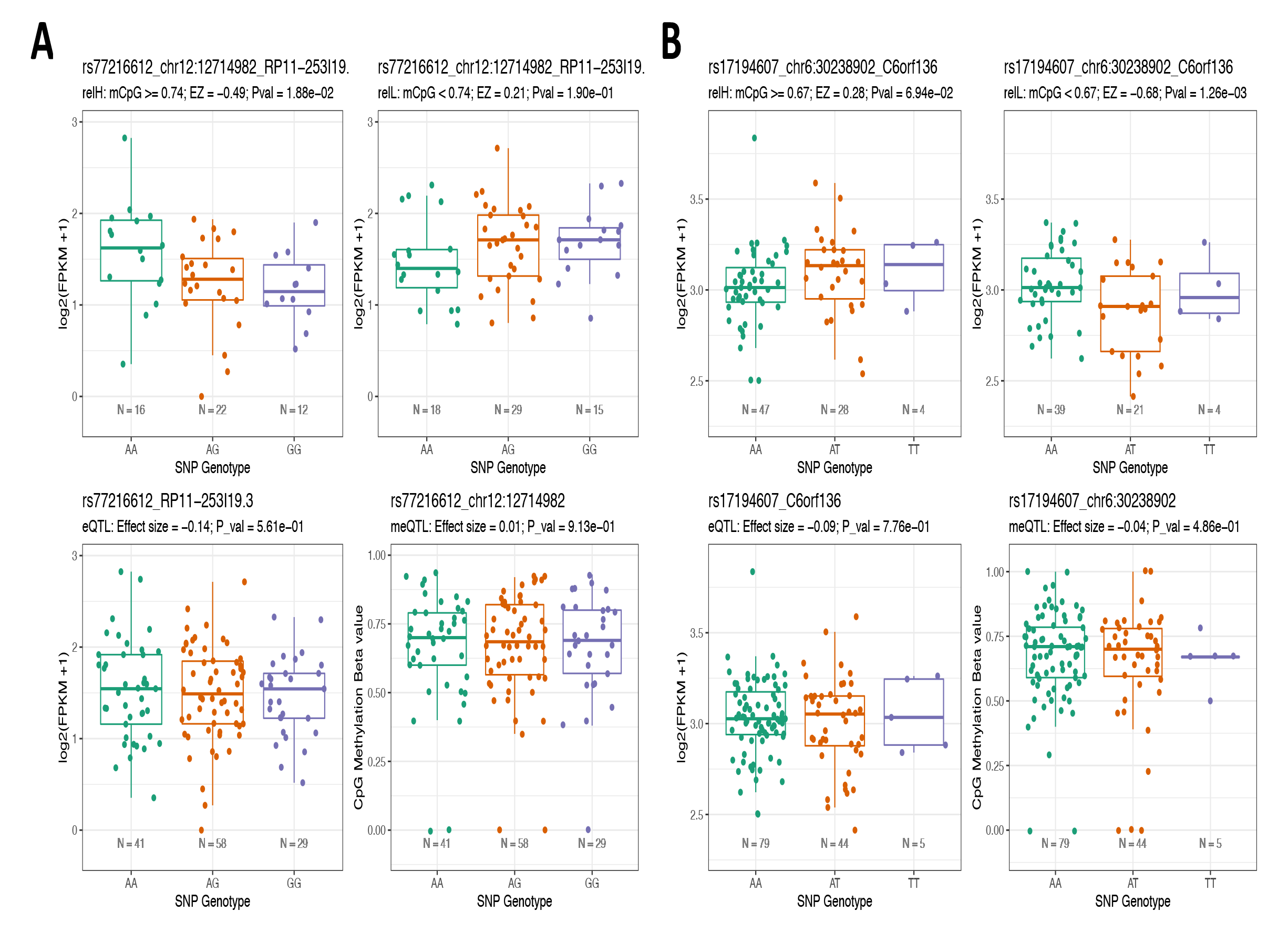
The memo-eQTLs help explain GWAS risk SNP. **A,B)** The visualization of memo-eQTLs, eQTLs and meQTLs for prostate cancer risk SNP rs77216612 and its high LD eSNP rs17194607.

**Supplementary Figure 4:**
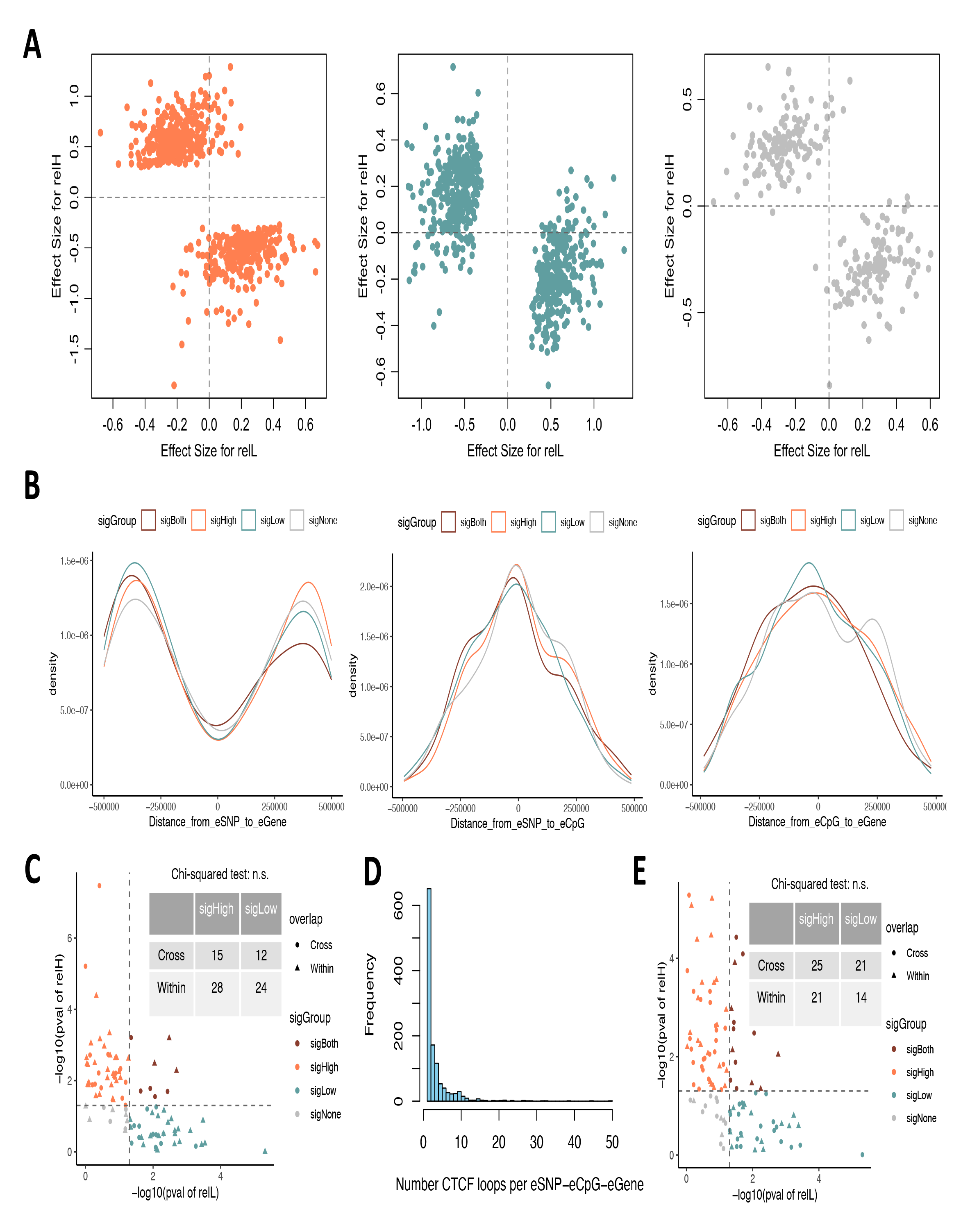
The memo-eQTLs help explain GWAS risk SNP. **A)** The effect size and direction of eSNPs on eGenes in eCpG relH and relL subsamples for the sigHigh, sigLow and sigNone memo-eQTLs. **B)** The distribution and comparison of distances between eSNP and eGene (left), eSNP and eCpG (middle) and eCpG and eGene (right) for the four memo-eQTL groups. Negative distances indicate the former element is on the left side of the latter element (Wilcoxon rank-sum two-sided test between sigHigh and sigLow groups: left: p = 6.08 × 10^-2^; middle: p = 6.97 × 10^-2^; left: p = 3.31 × 10^-1^). **C)** Overlapping patterns between eSNP-eCpG-eGene loci and eCpG-CTCF loops derived from RWPE1 ChIA-PET data for the four memo-eQTL groups (Chi-square test: p = 1). **D)** The number of VCaP HiChIP data derived CTCF loops that overlapped with the eSNP-eCpG-eGene loci. **E)** The overlapping patterns between the eSNP-eCpG-eGene loci and eCpG-CTCF loops derived from VCaP HiChIP data for the four memo-eQTL groups (Chi-square test: p = 7.78 × 10^-1^).

## Supplementary Tables

**Supplementary Table 1. Significantly correlated meCpG-CTCF pairs.** All significantly correlated meCpG-CTCF pairs (sheet1) and the most significantly correlated meCpG-CTCF pair per each CTCF binding site (sheet 2).

**Supplementary Table 2. memo-eQTL mapping results for 1,731 memo-eQTLs.**

**Supplementary Table 3. memo-eQTL eSNPs that overlap with GWAS risk SNPs.**

**Supplementary Table 4. Linear and spatial patterns for memo-eQTLs.** Pairwise linear distances between eSNP, eCpG and eGene for 1,711 memo-eQTLs (sheet1). The spatial overlapping patterns between the eSNP-eCpG-eGene loci and eCpG-CTCF loops derived from RWPE1 ChIA-PET data (sheet2) and VCaP HiChIP data (sheet3).

